## Supplementary Figures for "CASP microdomain formation requires cross cell wall stabilization of domains and non-cell autonomous action of LOTR1"

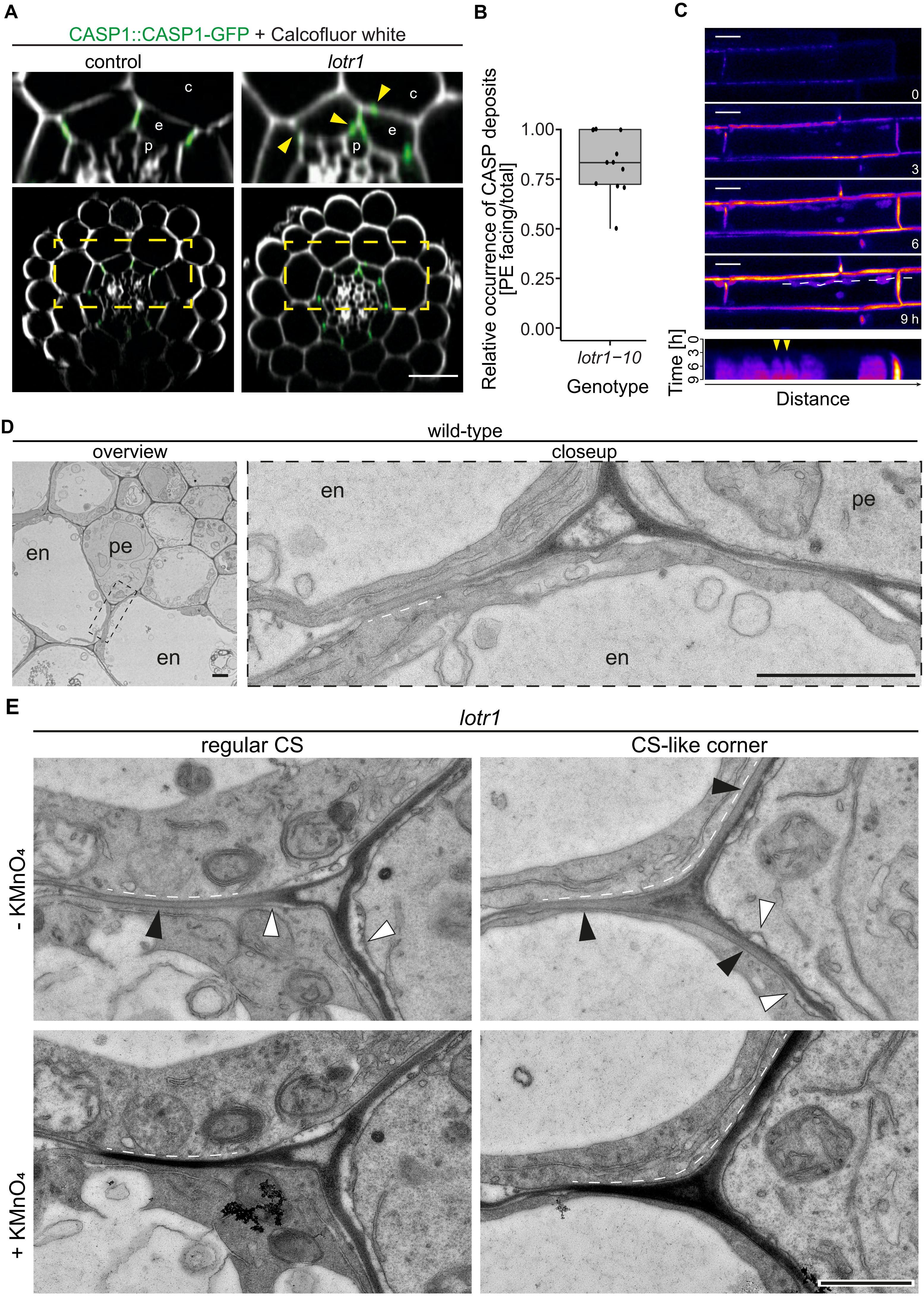
A
CASP1::CASP1-GFP + Calcofluor white
control
lotr1
c
e
p
B
Relative occurrence of CASP deposits [PE facing/total]
1.00
0.75
0.50
0.25
0.00
lotr1−10
Genotype
C
0
3
6
9 h
Time [h]
Distance
D
wild-type
overview
closeup
en
pe
E
lotr1
regular CS
CS-like corner
- KMnO₄
+ KMnO₄

Fig. S1

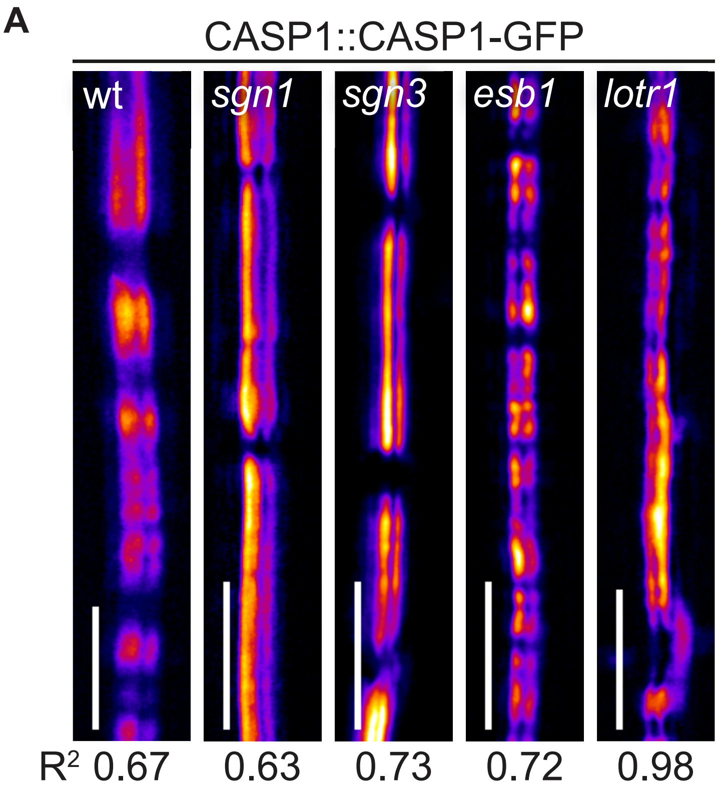
A
CASP1::CASP1-GFP
wt
sgn1
sgn3
esb1
lotr1
R2 0.67
0.63
0.73
0.72
0.98

**Fig. S2**

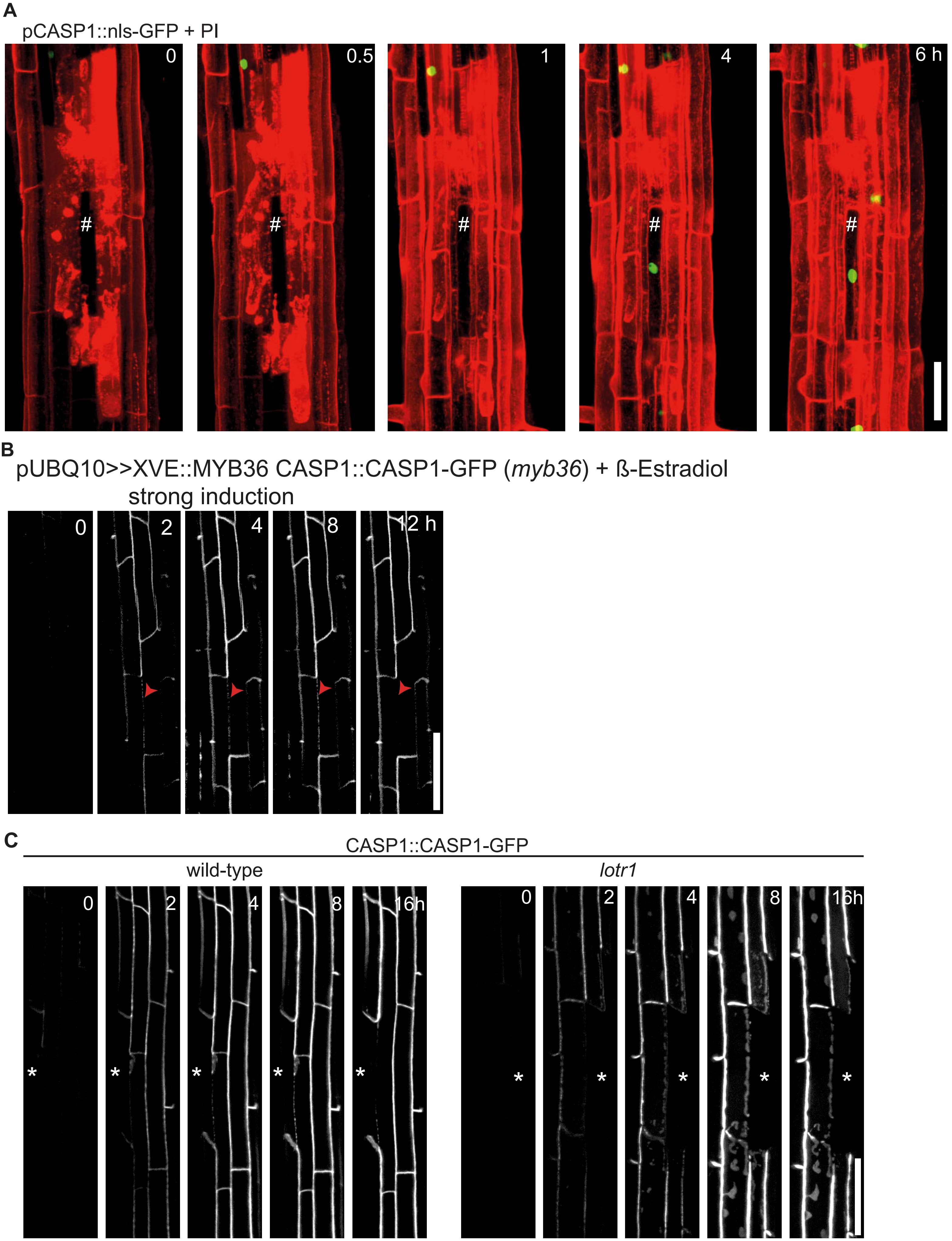
A
pCASP1::nls-GFP + PI
0
0.5
1
4
6 h
#
B
pUBQ10>>XVE::MYB36 CASP1::CASP1-GFP (myb36) + ß-Estradiol
strong induction
0
2
4
8
12 h
C
CASP1::CASP1-GFP
wild-type
lotr1
0
2
4
8
16h
*

**Fig. S3**

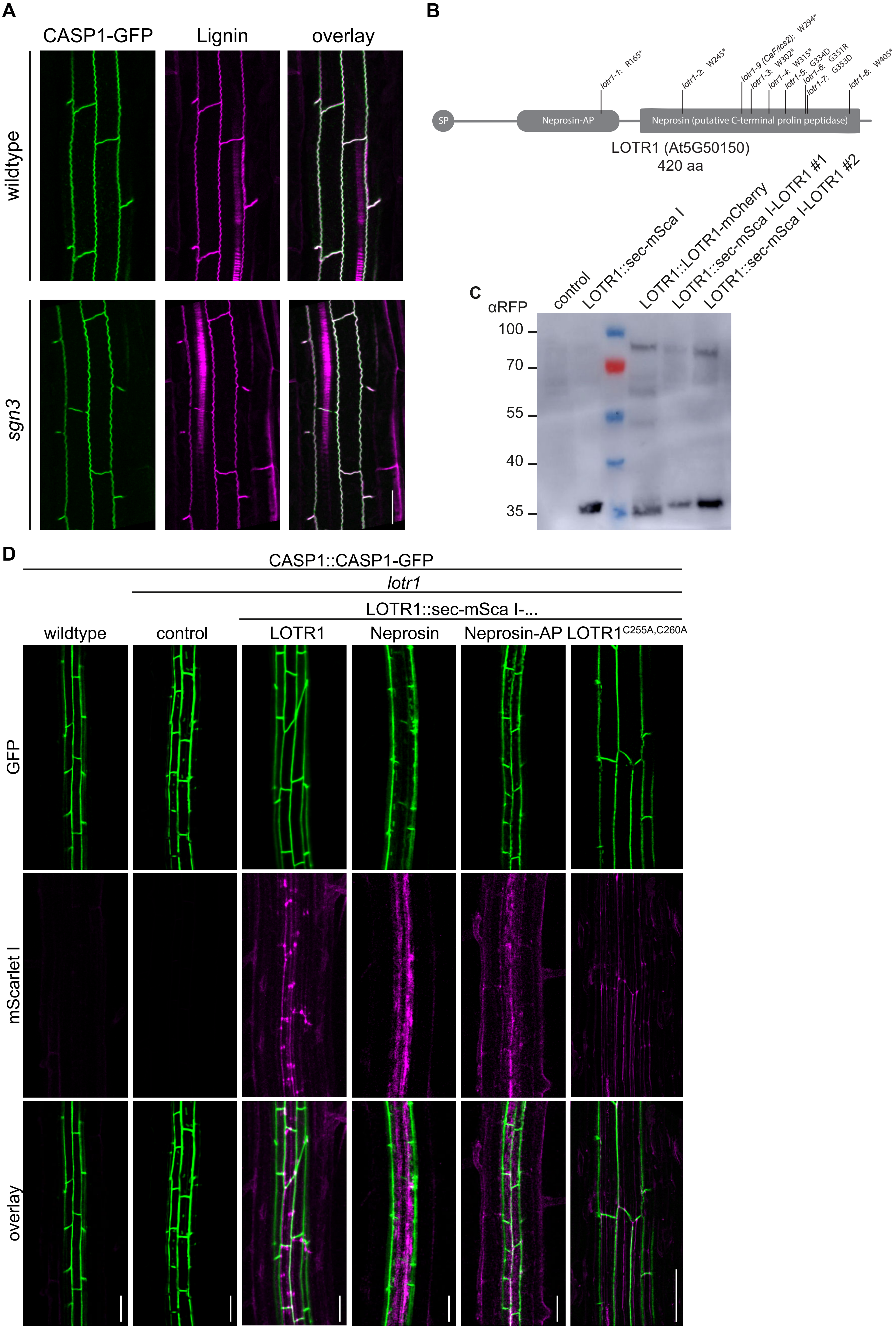
A
CASP1-GFP
Lignin
overlay
wildtype
sgn3
B
SP
Neprosin-AP
Neprosin (putative C-terminal prolin peptidase)
lotr1-1: R165*
lotr1-2: W245*
lotr1-9 (CaF/lcs2): W294*
lotr1-3: W302*
lotr1-4: W315*
lotr1-5: G334D
lotr1-6: G351R
lotr1-7: G353D
lotr1-8: W405*
LOTR1 (At5G50150)
420 aa
C
αRFP
control
LOTR1::sec-mSca I
LOTR1::LOTR1-mCherry
LOTR1::sec-mSca I-LOTR1 #1
LOTR1::sec-mSca I-LOTR1 #2
100
70
55
40
35
D
CASP1::CASP1-GFP
lotr1
LOTR1::sec-mSca I-...
wildtype
control
LOTR1
Neprosin
Neprosin-AP
LOTR1C255A,C260A
GFP
mScarlet I
overlay

**Fig. S4**

A

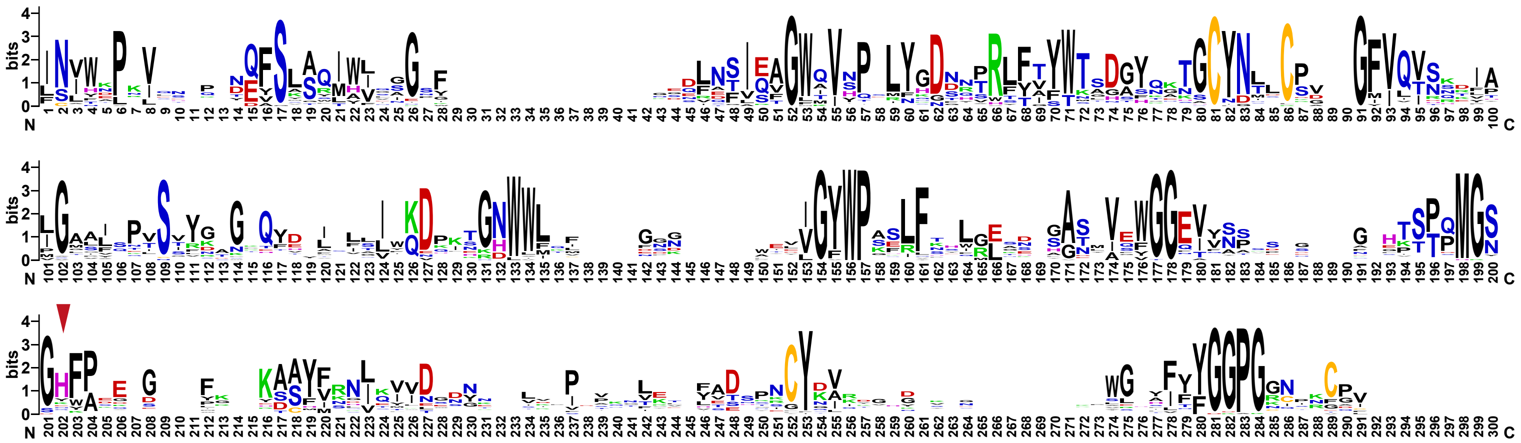

B

```
mepS (E.coli) CSANNTAKNMHPETRAV--GSETSSLQASQDEFENLVRNVDVKSRIM-DQYAD-------
     Neprosin QGANATINIWEPNLKNPNGDFSLTWIWISAGS-GSSLNTIEAGWQVYPGRTGDSQPRFFI
        LOTR1 ------INVWAPRVTDAY-EFSLSQIWLISGSFGHDLNTIEAGWQVSPELYGDNYPRFFT
    At1g10750 ------INVWAPQVQNQY-EFSLSQIWIISGSFGNDLNTIEAGWQVSPELYGDNYPRFFT
    At1g70550 ------INVWSPRVTSQY-EFGLSQIWVIAGSFTHDLNTIEAGWQISPELYGDTYPRFFT
    At1g23340 ------INVWTPRVISQY-EFSLSQIWIIAGSFAGDLNTIEAGWQISPELYGDTNPRFFT
                     :   *           : :.   ..    :..::.  .:      .*

mepS (E.coli) -WKGVRYRLGGSTKKGIDCSGFVQRTFREQFGLEL-PRSTYE--QQEMGKSVSR------
     Neprosin YWTADGYTSTGCYD--LTCPGFVQTNNYYAIGMALQP-SVYGGQQYELNESIQRDPATGN
        LOTR1 YWTTDAYQATGCYN--LLCSGFVQTNNKIAIGAAISPRSSYNGRQFDIGLMIWKDPKHGH
    At1g10750 YWTNDAYQATGCYN--LLCSGFVQTNSEIAIGAAISPSSSYKGGQFDITLLIWKDPKHGN
    At1g70550 YWTSDAYRTTGCYN--LLCSGFVQTNRRIAIGAAISPRSSYKGGQFDISLLIWKDPKHGH
    At1g23340 YWTSDAYQATGCYN--LLCSGFVQTNNRIAIGAAISPVSSYKGGQFDISLLIWKDPKHGH
               *.   *   *. .  : *.**** .    :*  : * * *   * ::   : .

mepS (E.coli) -------------------SNLRTGDLVLFRAG-------STGRHVGIYIGNNQFVHAST
     Neprosin WWLYL-WGTVVGYWPASIYNSITNGADTVEWGGEIYDSSGTGGFHTTTQMGSGHFPTEGY
        LOTR1 WWLELGNGLLVGYWPAFLFSHLRSHASMVQFGGEVVNSR-SSGAHTGTQMGSGHFADEGF
    At1g10750 WWLEFGSGILVGYWPSFLFTHLKEHASMVQYGGEIVNSS-PFGAHTSTQMGSGHFAEEGF
    At1g70550 WWLQFGSGALVGYWPAFLFTHLKQHGSMVQFGGEIVNNR-PGGSHTTTQMGSGHFAGEGF
    At1g23340 WWLQFGSGTLVGYWPVSLFTHLREHGNMVQFGGEIVNTR-PGGSHTSTQMGSGHFAGEGF
                                 . :      :  .*       . * *.   :*..:*   .

mepS (E.coli) SSGVIISSMNEPY-WKKRYNEA---RRVLSRS----------------------------
     Neprosin GKASYVRDLQCVDTYGNVISPTANSFQGIAPAPNCYNYQFQQGSSEL---YLFYGGPGCQ
        LOTR1 EKAAYFRNLQVVD-WDNNLLPLKN-LHVLADHPACYD--IRQGKNNVWGTYFYYGGP---
    At1g10750 TKSSYFRNIQVVD-WDNNLVPSPN-LRVLADHPNCYD--IQGGSNRAWGSYFYYGGP---
    At1g70550 GKASYFRNLQIVD-WDNTLIPASN-LKILADHPNCYD--IRGGTNRVWGNYFYYGGP---
    At1g23340 GKASYFRNLQMVD-WDNTLIPISN-LKVLADHPNCYD--IRGGVNRVWGNFFYYGGP---
               ..  . .::    : :         . ::
```

**Fig. S5**
